# Exploration of Anti-Yo and Anti-Hu paraneoplastic neurological disorders by PhIP-Seq reveals a highly restricted pattern of antibody epitopes

**DOI:** 10.1101/502187

**Authors:** Brian O’Donovan, Caleigh Mandel-Brehm, Sara E. Vazquez, Jamin Liu, Audrey V. Parent, Mark S. Anderson, Travis Kassimatis, Anastasia Zekeridou, Stephen L. Hauser, Sean J Pittock, Eric Chow, Michael R. Wilson, Joseph L. DeRisi

**Affiliations:** Department of Biochemistry and Biophysics, University of California, San Francisco, CA; UC Berkeley-UCSF Graduate Program in Bioengineering, University of California, Berkeley and San Francisco, CA; Diabetes Center, University of California, San Francisco, San Francisco, CA 94143; Department of Neurology, Department of Laboratory Medicine and Pathology, Mayo Clinic, Rochester MN; Weill Institute for Neurosciences, Department of Neurology, University of California, San Francisco, CA; Chan Zuckerberg Biohub, University of California, San Francisco, CA

## Abstract

Paraneoplastic neurological disorders (PNDs) are immune-mediated diseases of the nervous system understood to manifest as part of a misdirected anti-tumor immune response. Identifying PND-associated autoantibodies and their cognate antigens can assist with proper diagnosis and treatment while also enhancing our understanding of tumor-associated immune processes, triggers for autoimmune disease, and the functional significance of onconeuronal proteins. Here, we employed an enhanced version of phage display immunoprecipitation and sequencing (PhIP-Seq) leveraging a library of over 731,000 unique phage clones tiling across the entire human proteome to detect autoantibodies and create high-resolution epitope profiles in serum and CSF samples from patients suffering from two common PNDs, the anti-Yo (n = 36 patients) and anti-Hu syndromes (n = 44 patients). All patient samples positive for anti-Yo antibody by a validated clinical assay yielded polyspecific enrichment of phage presenting peptides from the canonical anti-Yo (CDR2 and CDR2L) antigens, while 38% of anti-Hu patients (17/44) had a serum and/or CSF sample that significantly enriched peptides deriving from the ELAVL family of proteins, the anti-Hu autoantigenic target. The anti-Hu antibodies showed a remarkably convergent antigenic signature across 15/17 patients corresponding to residues surrounding and including the degenerate motif, RLDxLL, shared by ELAVL2, 3 and 4. Lastly, PhIP-Seq identified several known and novel autoantigens in these same patient samples, representing potential biomarkers that could aid in the diagnosis and prognosis of PND and cancer.

## INTRODUCTION

Paraneoplastic neurological disorders (PNDs) affecting the central nervous system (CNS) are a collection of diseases characterized by an autoimmune response to proteins nominally restricted to the CNS and triggered by ectopic expression of the antigen by a systemic neoplasm.^1^ Neurologic symptoms of PNDs vary and can reflect damage to essentially any level of the neuroaxis. This symptom variability is, in part, determined by the neuroanatomic and/or cell type specificity of each PND. Existing PND pathogenesis models posit that an otherwise effective anti-tumor immune response to tumor antigens precipitates a breakdown of immune tolerance and subsequent infiltration of autoreactive lymphocytes and/or pathogenic autoantibodies into the CNS.^2^ The roles and culpability of cell-mediated and humoral immune responses in disease progression are disputed, as onconeuronal antibodies are often specific for intracellular proteins and are frequently detected in the sera of cancer patients in the absence of PND symptoms.^3^ In a typical clinical presentation, the neurological deficits and symptoms manifest before any underlying cancer is detected. As such, patients presenting with PND symptoms and tumor-associated antibodies are presumed to have cancer until otherwise ruled out and long-term tumor monitoring is recommended. Patients developing PND symptoms have lower rates of metastasis and higher rates of cancer survival, likely a consequence of the otherwise appropriate onco-immune response. Additionally, a majority of PND patients test positive for multiple tumor/PND related autoantigens.^4^ Antibody specificities are thought to be reflective of a tumor’s onconeural antigen expression, and clusters of neural autoantibodies in a patient’s serum or CSF can aid in specific cancer diagnosis.^5^

Anti-Yo paraneoplastic cerebellar degeneration (PCD) accounts for roughly 50% of subacute PCD presenting in middle age.^6^ The disease is marked by severe cerebellar ataxia, dysarthria, and Purkinje cell death and almost exclusively affects women, as it is primarily associated with gynecological and breast malignancies.^7^ Anti-Yo antibodies, also known as anti-Purkinje cell cytoplasmic antibody 1 (PCA-1), target the two intracellular proteins CDR2 (cerebellar degeneration related protein 2) and/or CDR2L (CDR2-like). Little is known about the function of these proteins other than that they have a role in transcription and that their expression is primarily limited to Purkinje neurons in the cerebellum, the brainstem, and spermatagonia.^8^ Whether the antibodies alone have pathogenic potential remains controversial; however, it is generally accepted that specific CD8+ T cells lead to Purkinje cell death.^9^ Although disease prognosis is poor, early detection, immunosuppression, and successful cancer treatment can limit Purkinje neuron loss and improve patient outcomes.^10^ To date, the specific antigenic determinants on the CDR2 and CDR2L proteins have not been mapped.

Anti-Hu-related PND is most commonly associated with small cell lung cancer (SCLC) and can present with differing neurologic syndromes ranging from dysautonomia, encephalopathy (limbic, brainstem, multifocal), epilepsy, cerebellar degeneration, myelopathy, and polyneuropathy.^11^ Anti-Hu antibodies target members of the ELAVL (embryonic lethal abnormal vision like) family of intracellular RNA-binding proteins. There are 4 members of this family, ELAVL1, ELAVL2, ELAVL3, and ELAVL4 (also known as HuA/R, HuB, HuC, and HuD, respectively). ELAVL1 is the most divergent homolog and is ubiquitously expressed in all tissues, while ELAVL2, 3, and 4 have restricted expression to the nervous system and are often referred to as nELAVL proteins (neuronal ELAVL).^12^

The majority of currently available clinical autoimmune panels include antigen-specific ELISA-based methods, cell-based assays (CBAs), co-localization immunohistochemistry (IHC), and traditional western blots.^13^ These candidate-based or single antigen tests may not capture the true complexity of the presumably more heterogeneous oncoantigenic repertoire. Unbiased and higher throughput methods exist in the research context and include immunoprecipitation-mass spectrometry (IP-MS), expression library screening, biochemical purification, peptide arrays, protein arrays, and phage display, each with its own advantages and limitations. Among these, variations on bacteriophage display technologies have been successfully utilized for decades in antigen discovery,^14^ epitope mapping,^15^ and antibody engineering.^16^ Traditionally, tissue-specific cDNA libraries or degenerate or random synthetic oligonucleotides are cloned and expressed on the surface of phage capsids which are panned against immobilized antibody or other targets of interest. More recently, rationally designed libraries encompassing the entire human proteome have been implemented.^17^ With next generation sequencing (NGS) as a readout, researchers can quantify the enrichment of millions of individual phage clones simultaneously and identify sequences that bind to the target or antibody of interest. This technology, known as Phage Immunoprecipitation and Sequencing (PhIP-Seq), enables high-throughput screening of patient samples and has successfully identified autoantigens and mapped epitopes in inclusion body myositis, rheumatoid arthritis, as well as two other common PNDs, the anti-Ma and anti-Ri syndromes.^18^ While broad spectrum autoantibody screening has previously been considered to be cost prohibitive,^19^ the continuing decline of large scale olignonucleotide synthesis and NGS costs may make whole proteome screening in by PhIP-Seq increasingly accessible in both research and clinical contexts.

Inspired by the potential of this technology to build upon our existing studies into the underlying causes of meningitis and encephalitis, we expanded on the published PhIP-Seq technique by designing a larger, more comprehensive library of endogenous human peptides. Our human “peptidome” consists of 731,724 unique phage clones and includes all known and predicted human protein splice variants. Using this expanded library, we conducted nearly 800 immunoprecipitations over 130 serum and CSF samples from patients suffering from anti-Hu and anti-Yo PNDs, revealing the most comprehensive and highest resolution epitope mapping of the autoantigens targeted in these PNDs to date. For the unusually convergent Anti-Hu epitope, additional deep-mutational scanning phage libraries where employed to map binding determinants at the single amino acid level. The results presented here further demonstrate the utility and high-resolution capability of programmable phage display for both basic science and clinical applications.

## RESULTS

### Design, Production, and Characterization of Human PhIP-Seq Library v2

The Human PhIP-Seq Library v2 contains all annotated human protein sequences from the NCBI protein database as of November 2015, including all published and computationally predicted splice variants and coding regions (Figure 1). Full-length sequences were clustered on 99% sequence identity (CD-HIT v4.6)^20^ to remove duplicate and partial entries, resulting in a set of 50,276 proteins. Each protein was computationally divided into 49 AA peptides using a 24 AA sliding window approach such that sequential peptides overlapped by 25-residues (see Methods). This resulted in a set of 1,256,684 peptide sequences encompassing the entire human proteome. To remove redundant sequences derived from identical regions of the various isoforms and homologs, we further clustered and collapsed the 49 AA peptides on 95% identity such that peptides with 2 or fewer AA differences were combined by choosing one representative sequence.

**FIGURE 1:**
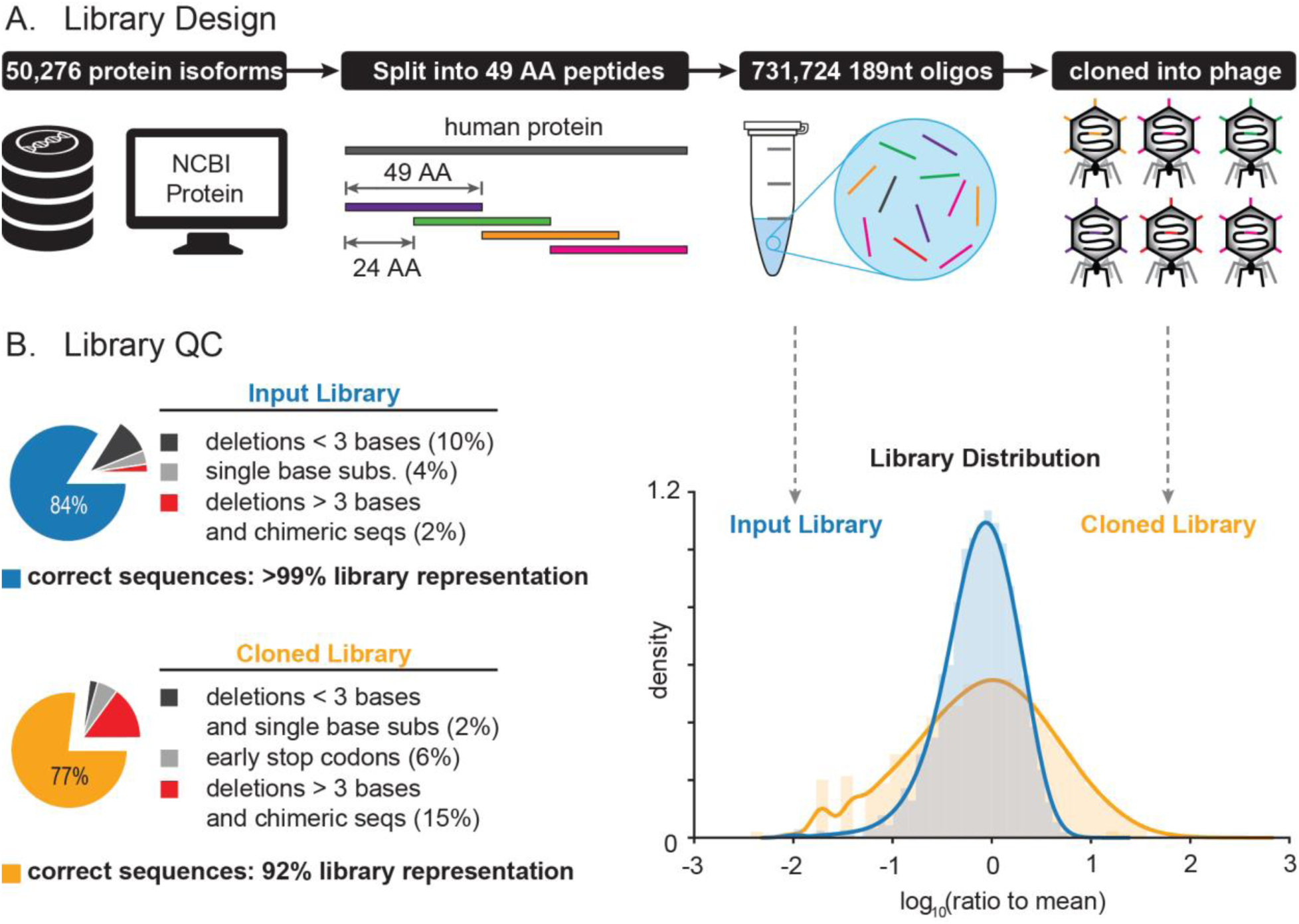
Design and characterization of PhIP-Seq Library. A.) All human protein isoforms, variants, and computationally predicted coding regions were downloaded from the NCBI Protein database. Full length sequences were clustered on 99% sequence identity (CD-HIT v4.6) to remove duplicate and partial sequences. Each protein was computationally divided into 49 amino acid peptides using a 24 AA sliding window. Redundant or duplicate sequences were removed by further clustering on 95% sequence identity. The resulting 731,724 sequences were synthesized and cloned into the T7 select vector (Millipore, Burlington, MA). B.) Assessment of the library quality before (blue) and after (yellow) cloning and packaging.

Amino acid sequences were converted to nucleotides using preferred *E. coli* codons (see Methods).^21^ Restriction sites (EcoRI, HindIII, BamHI, XhoI) were removed with synonymous nucleotide changes to facilitate cloning. To enable amplification from synthesized oligo pools, 21 nucleotide (nt) universal priming sites were added to the 5’ and 3’ ends of each sequence. These priming sites also encoded STREP and FLAG tag sequences (after addition of a terminal codon by PCR). Thus, the final library consisted of 731,724 unique 189mer oligonucleotide sequences encoding 49 AA human peptides flanked by 5’-STREP and 3’-FLAG sequences. The library was synthesized by Agilent Technologies (Santa Clara, CA) in three separate pools. The complete sequence of each oligo is available at https://github.com/derisilab-ucsf/PhIP-PND-2018.

After PCR amplification and size selection (Blue Pippin 3% Agarose cassette, Sage Science, Beverly, MA) but prior to cloning and packaging into phage, the synthesized oligos were evaluated by NGS with 70 million paired-end 125 nt reads (Figure 1B). 98.5% of all the synthesized sequences were full length, and 84% were error free. For those with errors, a majority (10% of all sequences) were deletions of 3 or fewer bases, while single base substitutions accounted for 4%, and the remaining 2% of oligos contained larger deletions (> 4 bases) or truncated/chimeric sequences. The commercially synthesized library yielded 722,436 unique peptide sequences with an estimated Chao1 diversity of 723,421, indicating that 99% of the library was present. The distribution and coverage of the library was uniform, with 99.9% of the sequences within ten-fold of the expected fraction (Figure 1C).

Size-selected and restriction-digested oligos were cloned and packaged in 25 separate 10 μl reactions according to the manufacturer’s specifications (EMD, Burlington, MA). The efficiency of *in vitro* packaging was quantified by plaque assay. The diluted and quenched packaging reaction (3 ml total) contained 10^9^ plaque forming units (pfu)/ml, roughly 1000x coverage of the entire library. The library of phage clones was sequenced (70 million paired-end 125 nt reads) to determine the fidelity of the packaged library. 77% of phage sequenced yielded error-free, full-sized inserts (Figure 1B). The majority of errors (21% of all sequences) were deletions or stop codons resulting in shorter expressed peptides. Within the 77% of correct sequences, there were 657,948 unique clones with an estimated Chao1 diversity of 658,476 – indicating that 89% of this library was packaged and cloned without any errors. Allowing for synonymous mutations and single amino acid substitutions, 92% of the library was packaged successfully into phage.

### Development of Statistical Approach to Analyze Peptide Enrichment

Several statistical approaches for analyzing PhIP-Seq data have been published, primarily based on Generalized Poisson (GP) or Negative Binomial (NB) count models fit to the distribution of unselected or input library phage populations.^22,23^ More recently, an algorithm leveraging z-scores of phage counts relative to AG bead only “mock” IPs was developed to account for non-specific phage binding to IP reagents and substrates.^24^ These approaches were developed to analyze datasets generated from single rounds of immunoprecipitation and were optimized for sensitivity to low abundance phage.

In order to identify phage significantly enriched after multiple consecutive rounds of selection with patient antibodies, a statistical framework was developed (see Methods) based on phage fold-change statistics relative to a collection of 40 independent mock IPs using protein-AG beads alone. Briefly, peptide counts from each control and sample IP were normalized to reads per 100,000 (rpK). Normalized peptide counts from each experimental IP were multiplied by a sample-specific scaling factor derived from the median rpK of the 100 most highly and consistently enriched AG-bead binding peptides (Supplemental Figure 1, sequences available at: https://github.com/derisilab-ucsf/PhIP-PND-2018). Each peptide’s fold-change enrichment was then calculated by dividing its scaled rpK value by its mean representation in the AG controls. Given the nature of the fold-change calculation, small fluctuations in low abundance phage in control IPs can lead to inflated fold-change values and false positives. To address this, fold-change values were further transformed using a method inspired by smoothed quantile normalization approaches in microarray data (see Methods).^25^ P-values were assigned to the fold-changes by evaluating the survival function of a normal distribution fit to the log10(fold-change) values in each experimental batch. Peptides reported as significant were those having an adjusted p-value (p_adj_) =< 0.05 in at least two replicates after multiple test correction (Benjamini-Hochberg).^26^

### Phage Library Performance Validation: Commercial Antibodies

Validation of the PhIP-Seq protocol was performed using two commercial antibodies with known specificities: anti-glial fibrillary acid protein (GFAP) (Agilent/DAKO, Carpinteria, CA) and anti-gephyrin (GPHN) (Abcam, Burlingame, CA) polyclonal antibodies. Three rounds of enrichment were used to further select for the particular epitope and reduce the proportion of non-specific binders. After each round of enrichment, phage populations were assessed by sequencing to an average depth of 2 million paired-end 125 nt reads. After 3 rounds of enrichment, a majority (50–70%) of the phage in the final population encoded peptides derived from the commercial antibody target (Figure 2A). Replicate IPs were conducted on separate days and exhibited high reproducibility (r > 0.8) in phage counts between experiments.

**FIGURE 2:**
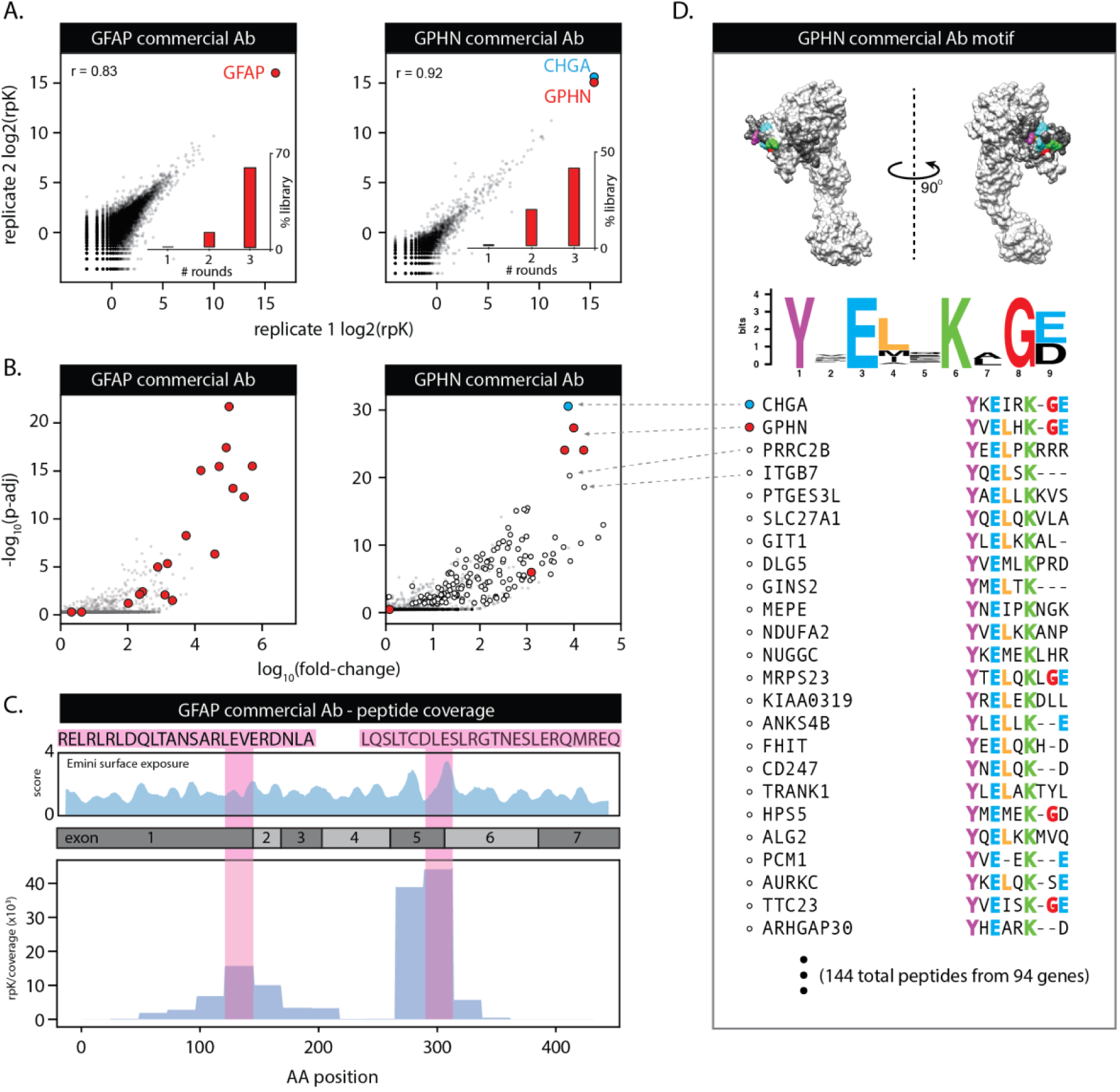
Validation of library by commercial antibody IP’s. Phage library was subjected to three rounds of selection by polyclonal commercial antibodies to GFAP and GPHN. A.) Replicates show high correlation of gene-level counts for IPs performed on separate days. Final libraries are dominated by the commercial antibody target. The GPHN antibody also consistently enriched a single peptide from chromogranin A (CHGA). B.) Scatterplots of peptide-level enrichments show 18 unique GFAP peptides and 4 GPHN peptides were identified as antibody binders (red markers). Peptides sharing a motif with both GPHN and CHGA (white, see D). C.) Alignment of GFAP peptides to the full-length protein reveals two distinct regions of antibody affinity corresponding to regions of high solvent exposure and B cell antigenicity as predicted by IEDB Tools Emini surface exposure and Bepipred algorithms. D). Motif analysis of the significant peptides in the GPHN IP reveal a short, discontiguous motif shared by 144 significantly enriched peptides in the final population (from 94 unique genes/proteins), including the most abundant CHGA peptide. The motif is highlighted on the crystal structure of GPHN (top) and peptides sharing this motif are marked by a white circle in B.

When applied to these data, our statistical model correctly identified peptides from the target gene of interest as the most significantly enriched with minimal off-target or unrelated peptides/genes (Figure 2B). The anti-GFAP IPs enriched 18 unique peptides representing > 60% of all phage in the final library (in both replicates) and scaled fold-changes greater than 300,000x (p_adj_ < 10^−20^) over the mock IPs. Similarly, the anti-GPHN commercial antibody enriched 4 unique C-terminal gephyrin peptides with 10,000-fold enrichment (p_adj_ < 10^−25^) over control IPs.

Alignment of the significantly enriched peptides from the anti-GFAP IP to the full-length protein revealed a distinct antigenic profile (Figure 2C), suggesting the commercial antibodies had highest affinity for a ~27 AA region surrounding the exon 5–6 junction. Notably, this region was identified by the immune Epitope Database (IEDB) tools Emini Surface Exposure calculation as having the highest B cell antigenicity potential.^27^ The three significantly enriched GPHN peptides were all overlapping sequences with consensus at the final 32 residues of the surface exposed C-terminus (Figure 1D). The anti-GPHN IPs also consistently enriched a single peptide from chromogranin A (CHGA) at levels greater than or commensurate with that of the GPHN peptides (p_adj_ < 10^−30^). Motif analysis (GLAM2)^28^ and alignment of the CHGA and GPHN phage-peptides, revealed a discontinuous 6-residue sequence (YxExxK) shared between CHGA and GPHN. Of the 256 peptides reported as significant in both replicates, 144 peptides (56%), contained the motif (Figure 2D). The probability of recovering these motif-containing peptides by chance given their representation in the original library is infinitesimal (< 10^−300^, Fischer’s exact test), indicating that a majority of the peptides in the final phage population were, in fact, true binders to the commercial antibody. These data also demonstrated that single AA resolution binding motifs could be recovered given sufficient representation in the original library.

### PND Cohort Results

In total, 798 individual IPs (including all three rounds of enrichment) were performed on 130 CSF and serum samples from patients with anti-Yo or anti-Hu antibodies identified by the Mayo Clinic Neuroimmunology Laboratory with CLIA and New York State approved methodology, as previously described (see Methods).^29^ The PhIP-Seq protocol was performed on a Biomek FX (Beckman Coulter, Brea, CA) automated liquid handler across three experimental batches (see Methods). All samples were run blinded and in duplicate. A sample was reported as positive if peptides derived from the respective Yo (CDR2/CDR2L) or Hu (ELAVL2,3,4) antigens_showed significant fold-changes (p_adj_ <= 0.05) in both replicates. Files containing the peptide counts for each IP are available at https://github.com/derisilab-ucsf/PhIP-PND-2018, and raw data are available under NCBI BioProject Accession PRJNA506115.

### Identification of CDR2/CDR2L antigens in anti-Yo sera and CSF

From 36 confirmed anti-Yo patients, a total of 53 samples were interrogated by PhIP-Seq, of which 51 (96%) were positive for anti-Yo peptides. When sample identities were unblinded, it was revealed that all 36 patients were represented by these positive samples. No significant differences were observed in sensitivity or binding profiles between CSF and serum, with 34/36 sera and 17/20 CSF testing positive (p_chi-square_ = 0.23). From 50 de-identified healthy donor sera, only a single CDR2L peptide from a single donor was significantly enriched (37 rpK, p_adj_ = 0.0494).

As shown in Figure 3A for three representative patients, phage specific for CDR2/CDR2L dominated the population by the final round of selection in a majority of samples. All patients showed enrichment for CDR2L while 30/36 (83%) were positive for both CDR2L and CDR2 peptides. No patients showed reactivity to CDR2 peptides alone. Across all patients, 29 unique CDR2L peptides were identified and accounted for 21% of all phage sequenced. Conversely, 6 unique peptides from CDR2 were identified across all patient samples and accounted for 0.25% of all phage sequenced (Figure 3B), suggesting that the overall magnitude and complexity of peptide enrichment for CDR2L was greater than for CDR2 peptides.

**FIGURE 3:**
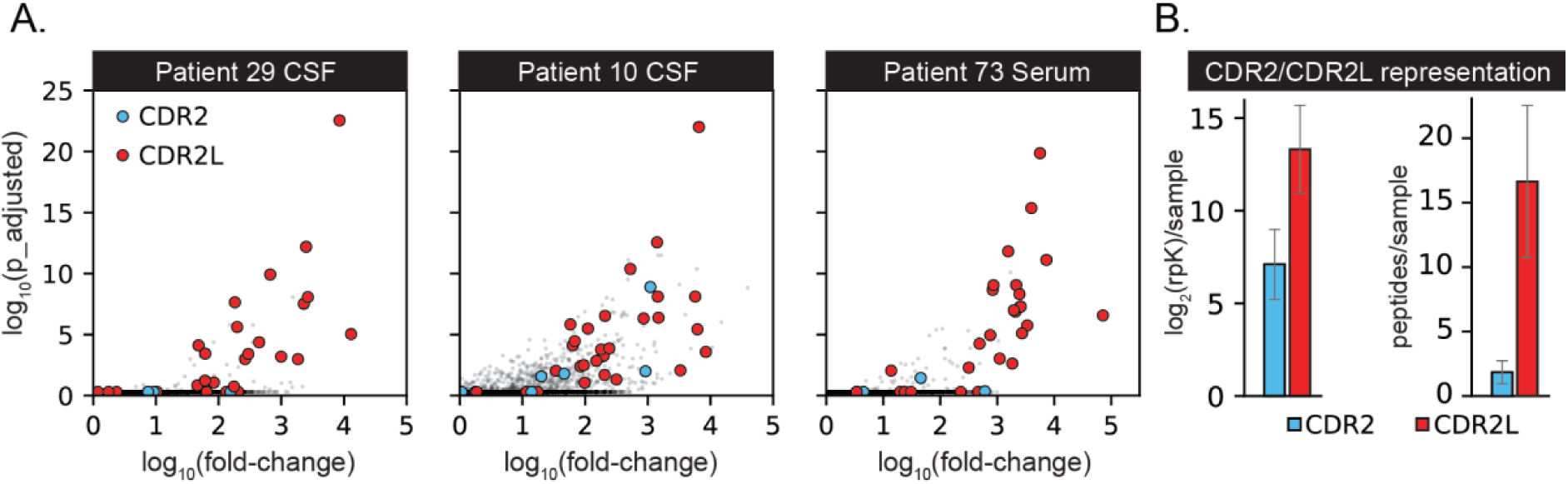
A.) Representative data from 3 patients demonstrating the robust enrichment of CDR2/CDR2L peptides. CDR2/CDR2L peptides were the top 5 most abundant peptides in 44/51 samples. B.) While CDR2/CDR2L peptides represented 21% of all phage, CDR2L enrichment is more pronounced and complex (more unique peptides per sample) than CDR2.

The antigenic profile across the full length CDR2L protein, for all 36 patients, is shown in Figure 4A. Convergent epitopes were shared by subgroups of patients, particularly at residues 322–346, the most highly enriched epitope across all patients. In contrast to CDR2L, the antigenic profile of patient antibodies to CDR2 (Figure 4C) was primarily limited to 3 peptides spanning the 100 AA at the N-terminus. We further-examined the relationship between homology and enrichment (Figure 4B). Interestingly, the most highly enriched regions of CDR2L represented the most divergent regions between the two proteins, suggesting that the antibody responses to CDR2L and CDR2 are independent and patient antibodies are, in fact, preferentially enriching for CDR2L over CDR2.

**FIGURE 4:**
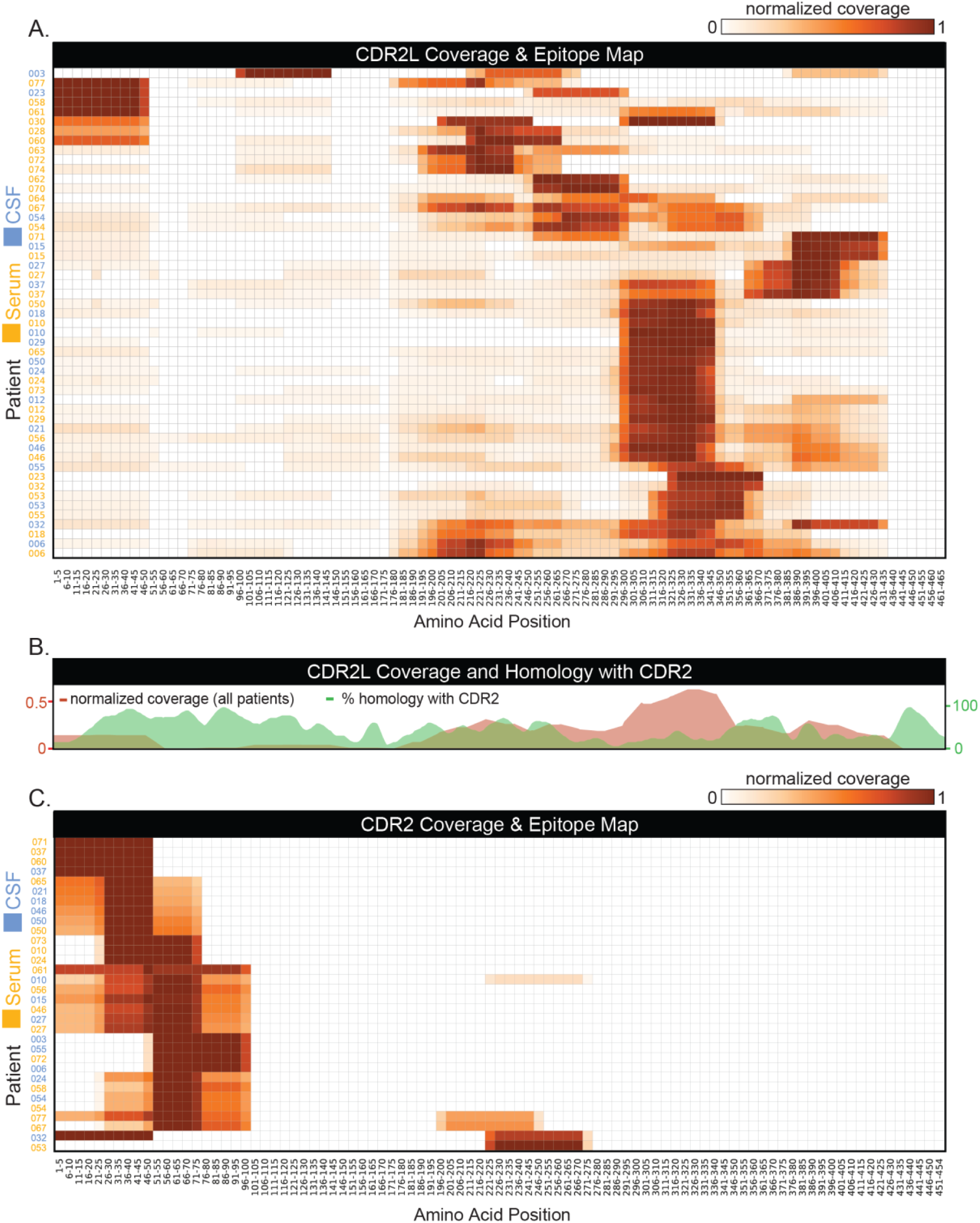
Anti-Yo cohort PhIP-Seq Results. A.) Epitope map showing normalized coverage of each samples’ significant CDR2L peptides aligned to the full-length gene (coverage is divided into 5 amino acid bins). Patient binding signatures vary, though a majority converge on residues 322–346. B.) Patient antibodies targeting CDR2L bind to regions of least homology with CDR2, suggesting the antibody responses to each protein are independent. C.) Epitope map showing normalized peptide coverage across the full length CDR2 sequence. Patient antibodies enrich for peptides primarily restricted to the first 100 amino acids at the N terminus. Abbreviations CDR2: cerebellar degeneration related protein 2, CDR2L: CDR2-like.

### Identification of nELAVL antigens in anti-Hu sera and CSF

From 44 patients with laboratory confirmed anti-Hu PND, a total of 76 samples (32 paired serum/CSF, two CSF, and 10 serum) were subjected to PhIP-seq. A total of 19 samples from 13 patients resulted in significantly enriched peptides (in both replicates) derived from the nELAVL genes (ELAVL2, ELAVL3, and ELAVL4), the known targets of anti-Hu antibodies. Across these 19 samples, 20 unique nELAVL peptides were identified. While the majority (>80%) of enriched peptides were attributed to ELAVL4 (HuD). For the purpose of visualization, all nELAVL peptides from each sample were aligned to the most common isoform of ELAVL4 (accession NP_001311142.1), shown in Figure 5A.

**FIGURE 5:**
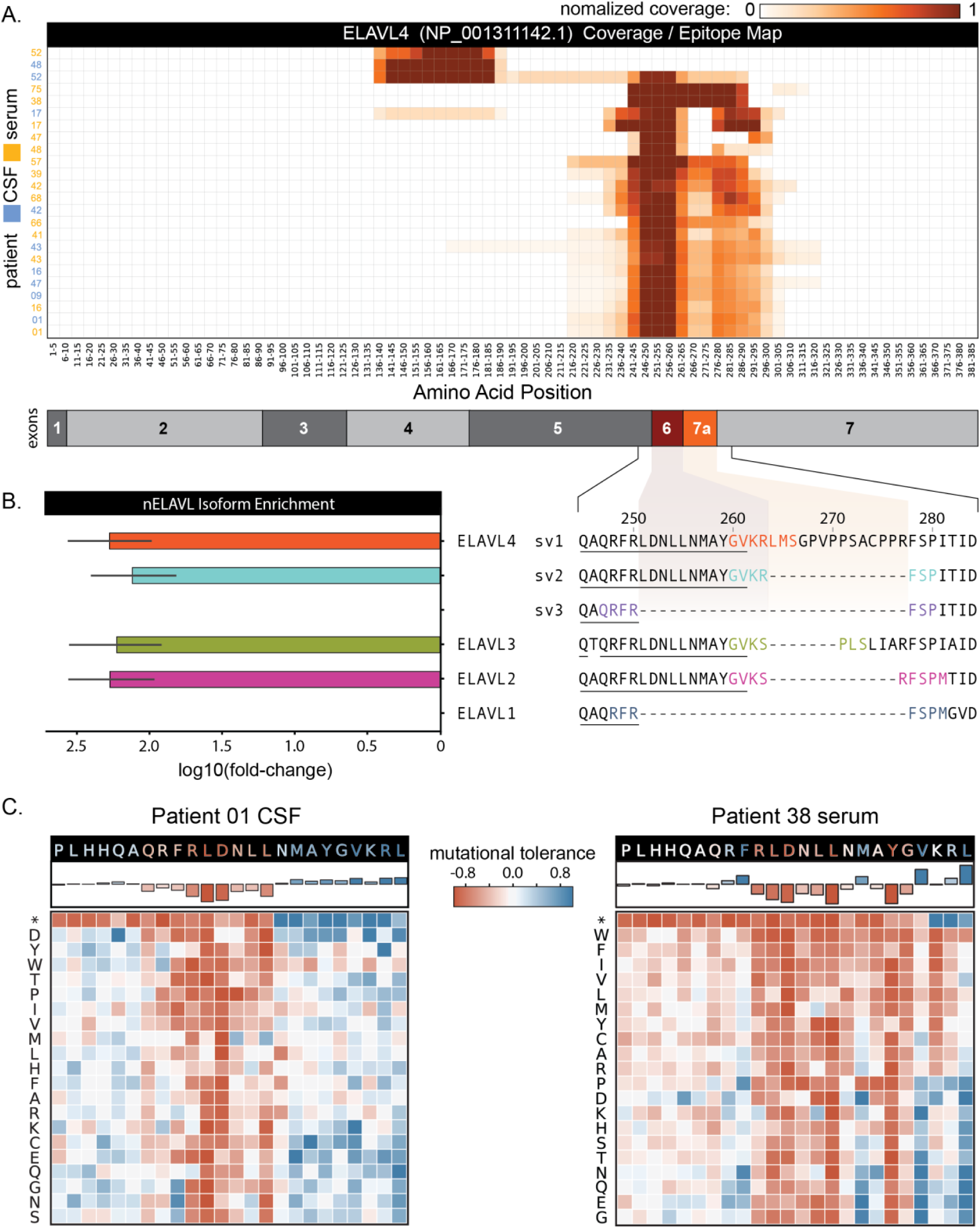
Anti-Hu PhIP-Seq Results. A.) Antibody binding signatures for samples that enriched nELAVL peptides. A majority of patients show a markedly convergent binding preference for a short 17-residue sequence at the exon 6/7a junction of ELAVL4. B.) Mean fold-change values for peptides spanning the unique regions defining ELAV4 splice variants and those from ELAVL2 and ELAVL3, indicating lack of enrichment beyond the exon 6/7a junction. C.) Mutational scan using smaller phage library with all possible point mutations at each location spanning the motif highlighted in (B). Mutational tolerance is calculated for each position, defined as the fold-change over unselected (AG-bead only IP) relative to the reference sequence. Most telling are the mutations encoding stop codons (denoted *) that reveal patient 01’s antibodies require a minimal sequence up to residues 255 (RLDDNLL) with the subsequence QRFRLDNLL least amenable to mutation. Patient 38’s antibodies bind to residues up to and including position 260 (RLDNLLNMAYGV), with highest affinity for RLDNLLN-AYG.

A majority (>90%) of the significantly enriched nELAVL peptides converged upon on a 17-residue sequence at AA positions 276–294 of ELAVL4, a sequence which is also common to variants of ELAVL2 and ELAVL3 (Figure 5B). The human peptidome PhIP-Seq library contained 18 peptides covering at least 10-residues of this short sequence, reflecting the large number of published isoforms in the NCBI Protein database for these highly spliced and studied proteins. Due to this redundancy, rpK and fold-change calculations were attenuated by as much as two orders of magnitude as individual peptide counts were distributed across multiple sequences containing the same epitope. To account for this, the analysis was re-run treating all 18 peptides containing this short sequence as a single entity, the anti-Hu “signature.” This resulted in the detection of an additional 8 samples from 4 patients yielding significant enrichment of the signature sequence. Notably, a single patient (patient 26) enriched only a single peptide in both serum and CSF derived from the N terminus of ELAVL2. The peptide is unique to ELAVL2 isoforms that had only been computationally predicted at the time of library design. These isoforms have since been confirmed experimentally and recognized as isoform c (accession NP_001338384.1).^30^ Importantly, none of 50 healthy, de-identified, patient sera samples yielded significant enrichment of nELAVL-derived peptides.

The convergent 17-residue signature sequence common to the majority of patient samples lies within a functionally significant hinge region between two RNA binding domains (RRM2 and RRM3) and may contain nuclear localization and export sequences responsible for control of subcellular localization and neuronal differentiation.^31^ Though exon and isoform naming conventions vary in the literature, this region is used to distinguish unique splice variants of ELAVL2, ELAVL3, and ELAVL4 (sv1, sv2, sv3) based on their inclusion or exclusion of relevant exons, specifically those commonly referred to as exons 6 and 7a. The identified motif terminates at the exon 6/7a junction (Figure 5B). The majority of patient samples preferentially enriched peptides that span the exon 6/7 junction, but do not include 7a, strongly suggesting that the binding determinants for the anti-Hu antibodies lie within this critical 17-residue anti-Hu signature region.

To further characterize the precise determinants of antibody binding, a new deep mutational scanning phage library was designed, encoding all possible single point mutants across the 17-residue signature motif (oligo design files available at https://github.com/derisilab-ucsf/PhIP-PND-2018). Samples from patients 01 and 38 were profiled by deep-mutational scanning PhIP-seq, as they represented the samples with the most and least complex peptide representation at this region, respectively. At each position, the fold-change enrichment of each amino acid substitution against an AG-only IP and relative to the abundance of the reference sequence is shown in Figure 5C, providing a landscape of the mutational tolerance at each residue. Examination of mutations that result in STOP codons (denoted by * in figure) are particularly informative, revealing the minimal sequence length required for antibody binding. Patient 01 antibodies required a minimal epitope containing residues up to and including positions 245–255 (QAQRFRLDNLL) with a shorter subsequence (QRFRLDNLL) being least tolerant of mutation. Patient 038’s antibodies appear to target a closely overlapping motif, RLDNLLN-AYG, with residues downstream not influencing peptide enrichment.

In light of this refined anti-Hu signature sequence, patient 01’s original PhIP-Seq data was re-analyzed using GLAM2 (Gapped Local Alignment of Motifs) to identify enriched motifs. Of the 36 peptides identified as significant in both replicates, 34 (94%) share a short, 7-residue motif, identical to the RLDxxLL identified by the mutational scanning approach. These include 16 peptides that were not derived from nELAVL genes, demonstrating that the dominant epitope in patient 01 is present in many unrelated protein sequences and explains the majority of peptides derived from non-nELAVL enriched genes. The biological significance of promiscuous binding to unrelated proteins harboring this motif is unknown.

### Consensus motif contains subsequences with predicted and experimentally demonstrated T-Cell Antigenicity

Examination of the refined anti-Hu signature motif and the full ELAVL4 sequence using the IEDB suite of sequence analysis tools (Supplemental Figure 2) suggests that the motif has low predicted probability of being a linear B cell epitope (Bepipred 2.0). The motif does, however, contain several sequences predicted to have high affinity for major histocompatibility complex type 1 (MHC-I) alleles (A1, A2, and A3 supertypes) and high prediction scores for MHC processing and presentation, with two predicted proteasomal cleavage sites flanking the region at amino acid positions 246 and 264. Interestingly, peptides derived from this region have previously been identified as anti-Hu T cell antigens.^32^ Indeed, peptides containing this exact motif have been shown to bind with high affinity to HLA-A1 (RLDNLLNMAY) and HLA-A2/B18 (NLLNMAYGV) and also to activate autologous CD8+ T cells obtained from Hu-positive patients.^33^

For paraneoplastic autoimmune diseases, the question of how immune tolerance is broken in the absence of new mutations remains to be addressed adequately. Selective exclusion of exons during expression in the thymus has been previously shown in mouse models as a possible route to break tolerance.^34^ In the case of the anti-Hu ELAVL family, splice variants have been previously shown to be controlled by a concentration-dependent autoregulatory mechanism wherein ELAVL proteins bind to AU-rich elements on nascent transcripts, protecting them from splicing and exclusion.^35^ Therefore, it may be possible that protein levels in the thymus during the development of central tolerance might fail to reach the requisite levels, resulting in exclusion of certain exons which in turn could affect processing and presentation. To further investigate this speculation, publicly available RNAseq datasets derived from bulk and sorted murine thymus and thymic medullary epithelial cells (mTECs) were examined. While most datasets lacked sufficient nELAVL coverage necessary for analysis of splice variants, a single mTEC from a single cell RNAseq dataset contained >50X coverage across ELAVL4, revealing the exclusion of exon 7a (Supplemental Figure 2).^36^ Comparatively, sorted murine hippocampal and cerebellar neurons show essentially even coverage across all exons.^37,38^

To further investigate exon inclusion and exclusion of nELAVL splice variants during the development of central tolerance, amplicon libraries spanning exons 5, 6,7a and 7 were generated from cDNA libraries derived from sorted human fetal and bulk adult thymic epithelial cells. An average of 1 million 125-nt paired-end reads (in triplicate) were obtained from each amplicon library. Exon 7a was represented in <0.5% of all reads in both fetal and adult TECs, supporting the notion that certain nELAV exons, such as exon 7a, are largely excluded from thymic expression. Interestingly, exon 7a is adjacent to exon 6 containing the critical anti-Hu signature motif. Taken together, these data point to a potential mechanistic connection to central thymic tolerance whereby incomplete T cell tolerance to selective exons within nELAVL helps promote an autoantibody response against this region of the protein.

### PhIP-Seq Identifies Additional Known and Potentially Novel Autoantigens in PND Samples

Antigens, other than those classically thought to be anti-Hu or anti-Yo were also investigated. Anti-CRMP5 (DPYSL5/CV-2) antibodies are a well-established clinical biomarker for SCLC and malignant thymoma and can be found together with anti-Hu antibodies in patients with a PND.^39^ Two of the anti-Hu patients (patients 51 and 52) had previously tested positive for anti-CRMP5 antibodies with a clinical assay (indirect immunofluorescence screening on mouse tissue substrate).^40^ This was corroborated by PhIP-Seq, with both patients’ CSF samples yielding significant enrichment of CRMP5 peptides. Patient 51 showed enrichment of 3 overlapping sequences at the C-terminus of the protein (AA residues 481–564) while patient 52’s CSF enriched for two peptides spanning residues 73–217. Though not contiguous on the primary sequence, these regions of the protein are in contact on the surface of the published three-dimensional structure (Figure 6).

**FIGURE 6:**
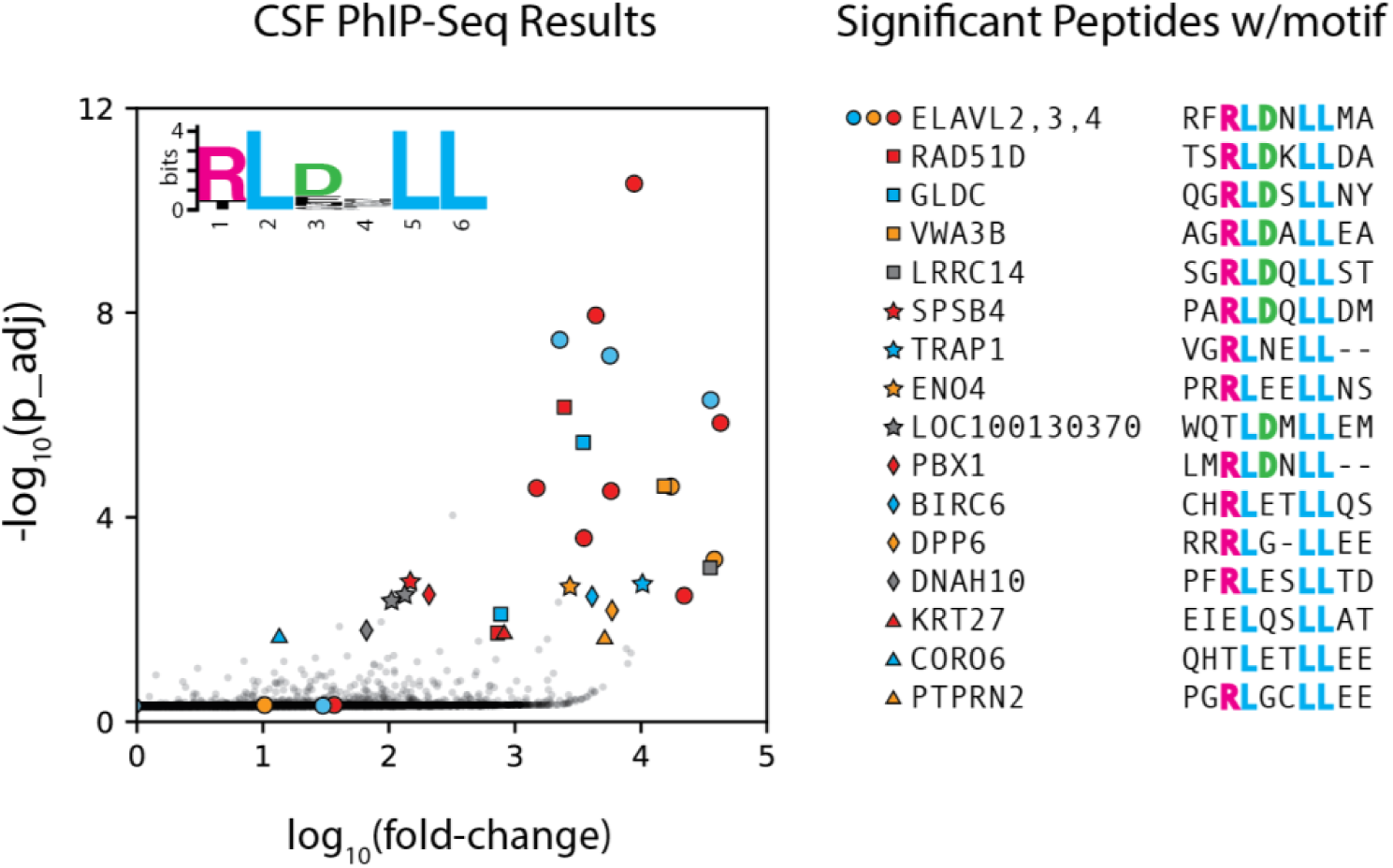
Immunodominant nELAVL motif in Patient 01 CSF sample is that identified by mutational scanning. The short sequence, dominated by RLDxLL is present in 34 of the 36 peptides identified in both replicates. Apart from nELAVL proteins, the patient’s antibodies enrich for peptides from 16 additional genes containing the motif. The biological consequences of such promiscuous autoantibody binding are unknown.

**FIGURE 7:**
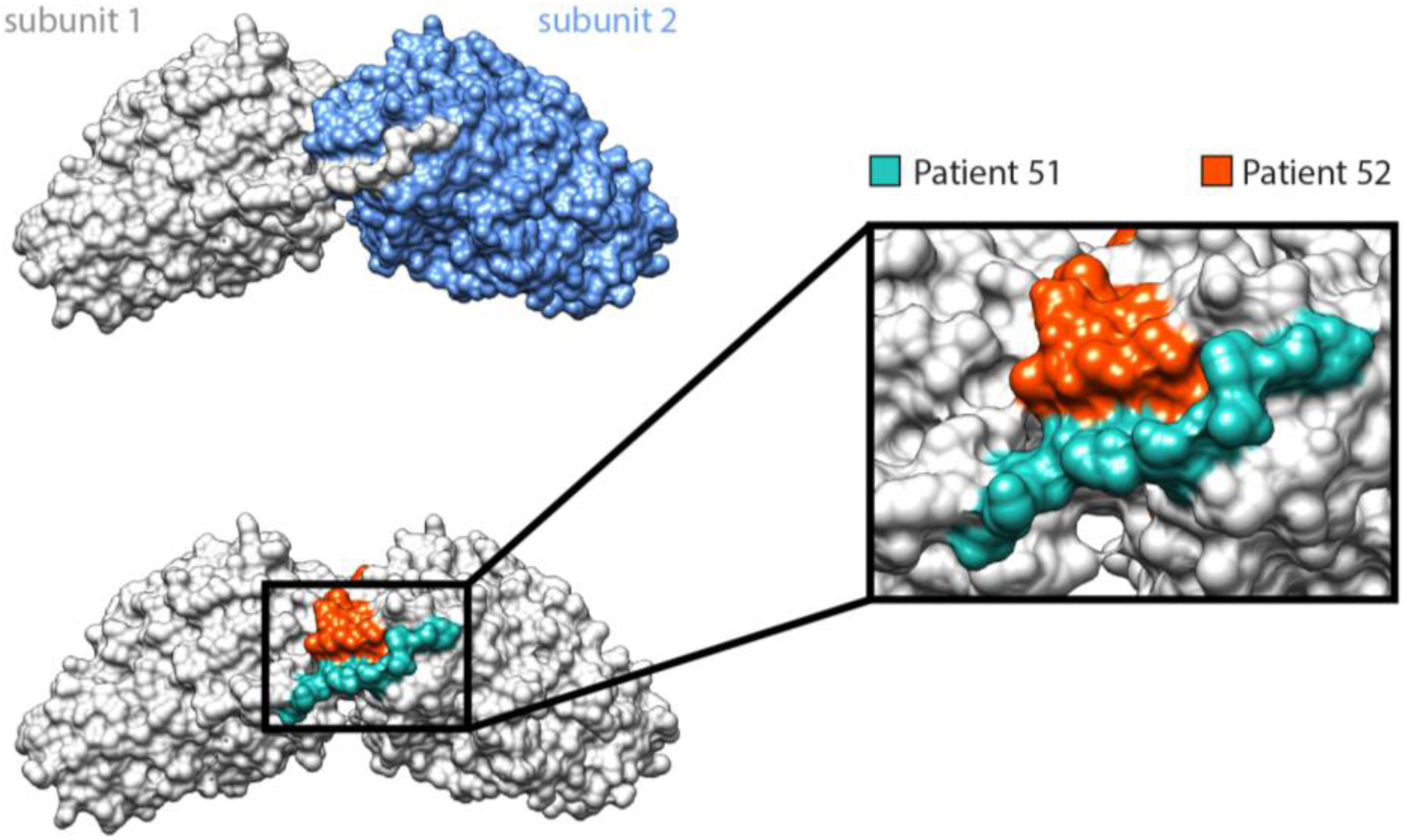
Anti-CRMP5 Antibody Epitope Mapping with PhIP-Seq. Two anti-Hu PND patient (patients 51 and 52) CSF samples also enriched CRMP5 peptides. CRMP5 is a well-established clinical biomarker for SCLC and malignant thymoma and anti-CRMP5 antibodies are often found concurrently with anti-Hu antibodies in patients with a PND. Though the peptides identified in each patient were discontiguous on the primary amino acid sequence, they are in close proximity in the 3D protein structure of CRMP5 at the region surrounding the C-terminus and the interface of the CRMP5 homodimer.

### Comparison to Healthy Sera Samples Identifies Disease-Associated Binding Signatures

To further detect additional disease-associated antigens, the PND cohort was compared to the binding signatures derived from the sera of 50 de-identified healthy subjects, thus allowing detection of known or potentially novel antigens by virtue of their enrichment at levels significantly higher than the healthy. To simplify analysis and avoid assumptions about the congruence of specific peptide binding signatures across multiple patients, we collapsed the data to the gene-level and called a patient sample positive for a given gene if it yielded enrichment of any peptides for a given gene in 2 or more samples/replicates. We compared the number of samples testing positive for each gene in the PND groups vs the control and determined significance by Fisher’s Exact test. The most significant gene-disease associations (FDR <0.05) are listed in Table 1. The full list of associations and healthy patient peptide counts are available at https://github.com/derisilab-ucsf/PhIP-PND-2018.

**TABLE 1:** Comparison of PhIP-Seq patient binding data to a collection of 50 healthy serum samples reveals known and potentially novel PND-associated gene-binding signatures. Genes enriched in PND samples (Fischer’s exact test, p-value <= 0.005). Genes showing significant associations after correcting for multiple tests (p_adj, Benjamini Hotchberg). SOX and ZIC family transcription factors are well established SCLC cancer biomarkers, and these antibodies are often associated with PND, particularly anti-Hu. In the anti-Yo cohort, PhIP-Seq enriched for PCM1, SAP25, ROBO4, and MTMR14.

| PND | Antigen | <u>Unique Peptides</u> |  | <u># samples</u> |  | **p** fisher's exact | **p** adj | Relevant Clinical Associations |
| --- | --- | --- | --- | --- | --- | --- | --- | --- |
|  |  | PND | Healthy | PND | Healthy |  |  |  |
| Hu | SOX family | 28 | 1 | 19 | 2 | $4.0 \times 10^{-06}$ | 0.01 | NSCLC, SCLC,<sup>41</sup> esophageal and bladder cancer,<sup>54</sup> |
| | ZIC family | 25 | 0 | 13 | 0 | $1.7 \times 10^{-05}$ | 0.01 | SCLC,<sup>55</sup> ovarian and breast cancer<sup>56,57</sup> |
| Yo | PCM1 | 7 | 0 | 24 | 0 | $2.1 \times 10^{-12}$ | $1.5 \times 10^{-9}$ | Parainfectious ataxia,<sup>43</sup> scleroderma,<sup>44</sup> glioblastoma<sup>58</sup> |
| | SAP25 | 16 | 0 | 15 | 0 | $3.6 \times 10^{-07}$ | $2.0 \times 10^{-4}$ | Ovarian cancer,<sup>59</sup> promyelocytic leukemia<sup>60</sup> |
| | ROBO4 | 4 | 1 | 15 | 1 | $4.1 \times 10^{-06}$ | $1.8 \times 10^{-3}$ | Breast cancer,<sup>61</sup> bladder cancer,<sup>62</sup> glioma<sup>63</sup> |
| | MTMR14 | 1 | 0 | 10 | 0 | $8.7 \times 10^{-05}$ | $3.2 \times 10^{-2}$ | none discovered |

### Zic and Sox Family Enrichment

Among proteins that were significantly enriched, many have been previously identified as serological markers of cancer. Antibodies to ZIC (Zinc Fingers of the Cerebellum) and SOX (SRY-related HMG-box) families of transcription factors have a well-established clinical association with SCLC and PND.^41,42^ Consistent with this fact, enrichment of peptides derived from the ZIC family (ZIC1, 2, 3, 4 and 5) was significant in 13/44 anti-Hu patients and 0/50 healthy controls (p_Fisher’s exact_ = 0.000017) while 19/44 patients and 2/50 healthy controls tested positive for SOX family peptides (p_Fisher’s exact_ = 0.000004). Though a majority of the peptides were derived from SOX1 and SOX2, the most common sequence, present in 13/19 patients with SOX reactivity, was a 34-residue sequence common to SOX1, 2, 3, 14, and 21. ZIC and SOX were the two most highly correlated genes in the anti-Hu patient data, with 7/13 patients with ZIC reactivity also enriching for a concurrent SOX peptide.

### Novel PND/Cancer Associated Signatures in Anti-Yo Patients

Beyond established PND and cancer-associated biomarkers, the data revealed several potentially novel disease-associated peptide binding signatures. After adjusting for multiple tests (Benjamini-Hochberg), PCM1 (Pericentriolar Material 1), SAP25 (Sin3A Associated Protein 25), ROBO4 (Roundabout Guidance Receptor 4), and MTMR14 (Myotubularin Related Protein 14) each show significant (p_adj_ <= 0.05) association with anti-Yo PND. Peptides derived from PCM1 showed the strongest association, with 24/36 anti-Yo patients and 0/50 healthy controls enriching a total of 7 unique peptides (p_Fisher’s exact_ < 10^−11^). PCM-1 antibodies have been reported in cases of parainfectious acute cerebellar ataxia and autoimmune scleroderma, ^43,44^ but this is, to our knowledge, the first reported case of anti-PCM1 activity associated with Anti-Yo PND. Multiple peptides from SAP25 and ROBO4 were each identified in 15 patients (5 patients concurrently) while a single peptide from MTMR14 was common to 10 anti-Yo patients and zero healthy control sera.

## DISCUSSION

We have designed and implemented the most complex and comprehensive human PhIP-Seq library to date, encompassing the entire human proteome and including all known and predicted splice variants and isoforms in the NCBI Protein database. In an effort to better understand autoimmune neurological diseases, we screened patient CSF and serum samples from two common paraneoplastic neurological disorders, the anti-Yo and anti-Hu syndromes. We chose these diseases for several reasons. First, while their autoantigens have been previously identified, the high-resolution epitope mapping afforded by PhIP-Seq or similar methods has, to our knowledge, not yet been achieved. Second, we reasoned that the high degree of alternative splicing in these neuronal antigens would help us leverage the unique design of our library which includes all published and computationally predicted splice variants. Third, as these diseases are associated with a robust anti-tumor immune response, we reasoned that patient samples would likely enrich for multiple autoantigens that could have disease relevance and would be missed by more limited and hypothesis-driven antibody assays.

To our knowledge, the PhIP-Seq data from the anti-Yo patient cohort represent the highest-resolution epitope mapping of anti-Yo antigens to date. PhIP-Seq correctly identified the canonical anti-Yo antigens, CDR2 and CDR2L in 36/36 patients. This highly concordant result suggests that anti-Yo PND antibodies routinely bind linear epitopes without any requisite post-translational modifications (PTMs) or significant secondary or tertiary structure. In addition, CDR2L peptides were much more enriched than CDR2 peptides. Indeed, across all samples, phage expressing CDR2L peptides represented 21% of all phage sequenced, and these peptides were represented in the top 5 most significantly enriched sequences in > 80% of samples tested. While these data do not provide a quantitative measurement of antibody affinity (especially to the native form *in vivo*), they suggest that CD2RL is the immunodominant antigen in patients suffering from anti-Yo PND. This also supports previous studies arguing that the antibody responses to each protein are independent (i.e., not cross-reactive) and that CDR2L is the primary antigen.^45^

Within the anti-Hu cohort, PhIP-Seq yielded significant enrichment of nELAVL peptides in 17/34 patients. Those patients that were positive for nELAVL peptides yielded a remarkably convergent antigenic signature, with antibodies binding a short 17-residue sequence (QAQRFRLDNLLNMAYGVK) shared by ELAVL2, 3 and 4 but absent in ELAVL1. This short sequence lies at the junction of exons 6 and 7a in ELAVL4, and high resolution deep mutational scanning followed by motif analysis led to a further refined signature motif sequence (RLDNLLNMAY) as most deterministic for antibody binding. We note that this short sequence has been previously characterized as a T cell epitope in anti-Hu patients.^32,33^ Regarding the anti-Hu patients that did not yield peptides from nELAVL genes, it may be the case that the autoimmune antibodies in these patients may require specific secondary and tertiary conformations, somatic mutations, or PTMs not emulated by our peptide library.

Interestingly, our analysis of publicly available RNAseq datasets and our own amplicon sequencing of human thymic epithelial cells reveals that exon 7a, immediately adjacent to our identified motif, is absent in >99% of ELAVL4 transcripts in the thymus but abundant in specific neuronal subtypes and brain regions in mice. While investigation of exon exclusion in the thymus and the mechanisms underpinning autoimmunity are beyond the scope of this initial survey, we speculate that the absence of these short exons during the development of central tolerance could potentially influence the rise of autoreactive T lymphocytes when the full-length protein is encountered in the CNS or expressed by peripheral tumors. Further, exclusion of exons bearing proteasomal cleavage sites (as with exon 7a) could bias MHC processing and presentation. Lineages of B cells with B cell receptors specific for ELAVL4 isoforms containing these excluded exons could presumably be maintained and expanded through interactions with CD4+ T cells specific for this short sequence when displayed on the surface of a B cell acting in its capacity as an antigen presenting cell. Such a collaborative process and mechanism is in keeping with previous studies examining autoimmune dymelinating disease and Nova2 PND.^34,46^

A major advantage of unbiased proteomic approaches like PhIP-Seq is the ability to interrogate patient samples for multiple antigens simultaneously, enabling high-throughput antigen-antigen and antigen-disease associations. By comparing the PND patient data with a collection of 50 healthy sera run on our platform, we identified several known and potentially novel antigens and/or biomarkers associated with each disease; most notably, ZIC and SOX family genes in the anti-Hu patients and PCM-1 and SAP25 in the anti-Yo cohort. The biological and clinical significance of these novel associations remain unknown. The clinical data on the specific cancer diagnosis and detailed neurologic phenotyping for each patient was not available for this study, and thus precludes analyses contrasting tumor type and particular neurologic deficits with the patient-level peptide binding signatures. Much like transcriptomic analyses, specific sets of correlated or biologically related antigens may provide researchers and clinicians with insight into yet undiscovered or previously unmeasurable cellular processes. We speculate, for example, that antibody binding signatures to endogenous, intracellular and normally inaccessible antigens reflect an immune process related to clearance of intracellular debris following apoptosis of tumor cells during the onco-immune response.^47^ The preponderance of ZIC and SOX related peptides in our anti-Hu cohort and their high levels of expression in SCLC, for example, supports the hypothesis that patient antibodies are providing a snapshot of the interior of tumor cells.^5^ The potential value of such biomarkers may only be realized after hundreds or thousands of patient samples have been run and correlated with higher-resolution disease phenotypic datasets that include long-term clinical outcomes and tumor transcriptional profiling.

This validation study confirms and extends the comprehensive nature of PhIP-Seq peptidome libraries for the purpose of identifying novel autoantigens in patient groups. Ultimately, as comprehensive, proteome-wide serological screens are more widely adopted and replace conventional candidate-based approaches, we expect the diagnostic, prognostic, and research potential of such assays to be fully realized. Indeed, the third most common category of autoantibody-mediated causes of encephalitis belonged to patients with unidentified autoantigens that have defied identification by more conventional methods. ^48^ Furthermore, high-resolution datasets characterizing the co-occurrence of specific antigens and antigenic determinants to specific diseases, symptoms, and clinical outcomes may ultimately serve to identify early biomarkers and may even stratify patient outcomes and inform treatment decisions. Finally, higher resolution epitope mapping and analysis of antigenic signatures across multiple patients should better inform the currently incomplete models describing onco-immunologic processes and the loss of self-tolerance underpinning the rise of PND and autoimmune disease in general.

## MATERIALS AND METHODS

### Computational Design of Library

We downloaded all human sequences in the NCBI protein database (Nov 2015); including all splicing isoforms and computationally predicted coding regions. Full length sequences were clustered on 99% sequence identity (CD-HIT v4.6)^20^ to remove duplicate and partial sequences before computationally dividing them into 49 amino acid peptides using a 24 AA sliding window approach starting at the N terminus. As such, each peptide shared a 25-residue overlap with the one preceding it on the primary protein sequence. Peptides encoding the final 49-residues at the c-terminus were substituted when the final sliding window resulted in shortened or truncated sequences (proteins not evenly divisible by 49). The resulting peptides were further clustered/collapsed on 95% identity such that such that peptides with 2 or fewer amino acid differences were combined by randomly choosing one representative sequence.

Amino acids sequences were converted to nucleotides using preferred E. *coli* codons (Supplemental Table 1) and relevant restriction sites (EcoRI, HindIII, BamHI, XhoI) removed with synonymous mutations to facilitate cloning. To enable amplification from synthesized oligo pools, 21 nt universal priming sites were added to the 5’ and 3’ ends of each oligo. These sequences also serve to encode STREP and FLAG tag sequences (after addition of terminal codons by PCR) that allow for downstream assessment of proper, in frame cloning. The resulting library consists of 731,724 unique 189mer oligonucleotide sequences encoding 49AA human peptides flanked by 8AA STREP and FLAG sequences after amplification with appropriate primers.

### Cloning into T7 Select vector

Single-stranded DNA oligonucleotides were commercially synthesized in three separate pools by Agilent Technologies (Santa Clara, CA). Oligos were amplified by PCR primers adding relevant EcoRI and HindIII restriction sites. 200 pg of DNA (10^8^ molecules per reaction across 50 reactions) was used as a template for each PCR reaction:

| Reagent | Vol. (μl) | Concentration |
| --- | --- | --- |
| Input DNA library | 1 | 200 pg/μl |
| Platinum HiFi master mix | 10 | 5X |
| Forward Primer*<br>(adds EcoRI site) | 2 | 10 μM |
| Reverse Primer*<br>(adds HindIII site) | 2 | 10 μM |
| H<sub>2</sub>O | 35 | - |

\*primer design:

Thermocycling Conditions:

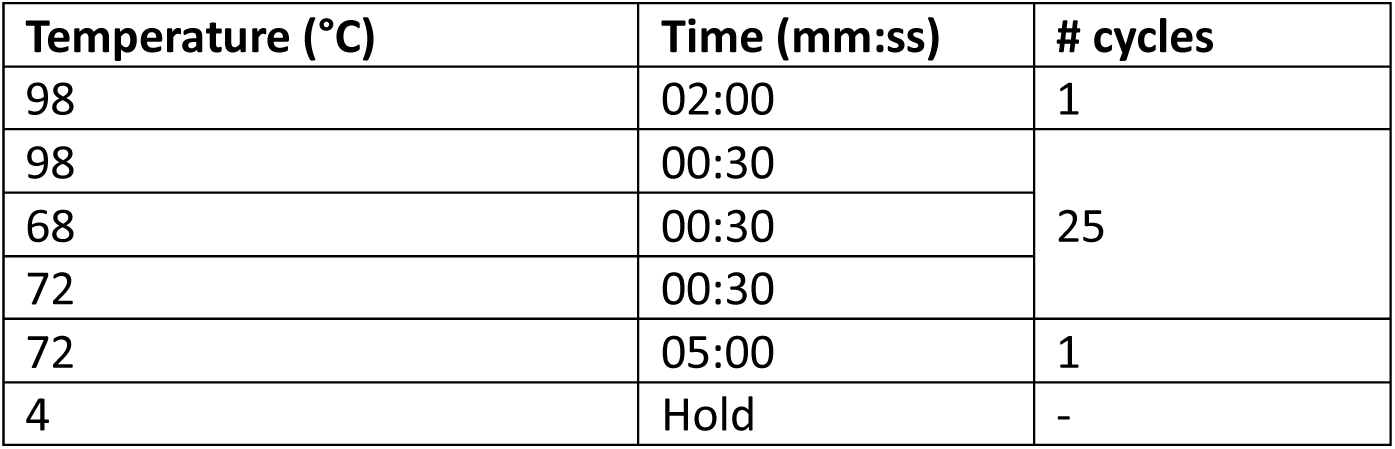

PCR products were column cleaned (Zymo DNA clean and concentrator, Zymo Research, Irvine, CA) and eluted in 20 μl H_2_O. DNA was then subject to restriction digestion under the following reaction conditions (incubated at 37 for 1 hour):

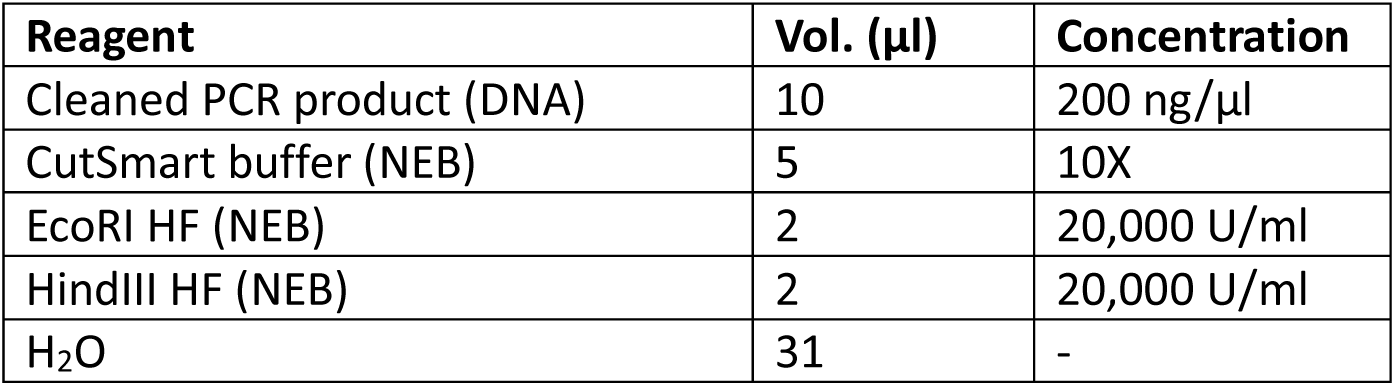

To minimize the proportion of truncations and deletion errors most commonly associated with commercial synthesis chemistries, digested fragments were size selected by agarose gel electrophoresis (Blue pippin 3% cassette - BDQ3010, Sage Sciences, Beverly, MA) before ligation/cloning.

### Cloning and Packaging

The digested oligonucleotides were cloned and packaged into competent phage following the T7 select cloning manual (EMD/Novagen). The following reaction mix was incubated at room temperature for 15 minutes:

| Reagent | Vol. (μl) | Concentration |
| --- | --- | --- |
| Digested, size-selected DNA | 1 | 6 ng/μl |
| T7 Vector Arms (EMD) | 3 | 500 ng/μl |
| Quick Ligase Buffer (NEB) | 5 | 2X |
| Quick Ligase (NEB) | 1 | 10,000 U/ml |

2 μl of the ligation reaction were added directly to 10 μl of T7 packaging extract (EMD) and allowed to incubate at room temperature for 2 hours. Reaction was quenched by adding 200 μl of chilled, sterile, LB. 24 separate 10 μl packaging reactions were carried out to ensure library complexity. Packaging efficiency was determined by plaque assay (as described in T7 Select Cloning Manual).

### In vivo amplification

BLT5403 E. *coli* cultures were grown to log phase (OD600 = 0.5) and inoculated (0.001 MOI) with packaging reaction and allowed to clarify (complete lysis) by incubating in 37 °C incubator (2–3 hours). BLT5403 carries an ampicillin-resistant plasmid expressing a wildtype gene 10A behind a T7 promoter. Phage produced in this strain carry 5–15 copies of the 10B capsid protein bearing the cloned fusion peptide. Lysates were cleared of debris by centrifugation at 4,000 rcf at 4 °C for 30 min before filtering through a 0.22 micron filter.

### Concentrating and Storing Phage Library

Phage were precipitated by adding 1/5 volume 5x PEG/NaCl precipitation buffer (PEG-8000 20%, NaCl 2.5 M) and centrifuged at 13,000g for 60 minutes. Pellets were raised in 1/4 starting volume storage buffer (20 mM Tris-HCl, pH 7.5, 100 mM NaCl, 6mM MgCl_2_).^49^ Stocks were titered by plaque assay and diluted/concentrated to ~10^11^ pfu/ml.

### Immunoprecipitation

All liquid handling steps were carried out on Biomek FX robotics platform (Beckman Coulter, Brea, CA). 96-well, full skirted, low profile PCR plates (BioRad Inc, Hercules, CA) were incubated overnight with 180 μl IP blocking buffer (3% BSA, PBS-T) to prevent nonspecific binding. Blocking buffer was replaced with 150 μl of phage library (10^11^ pfu in storage buffer) and mixed with 2 μl patients CSF, patient sera diluted 1:50 in blocking buffer, or 2 ng commercial antibody and incubated O/N at 4°C.

Protein A and G magnetic beads (Dynabeads – Invitrogen, Carlsbad, CA) were mixed equally, washed three times by magnetic separation and resuspension in equivalent volumes of TNP-40 wash buffer (150 mM NaCl, 50 mM Tris-HCL, 0.1% NP-40, pH 7.5). 10 μl of A/G bead slurry was added to each IP reaction (using wide bore pipette tips) and incubated for 1 hour at 4°C. Bead-antibody complexes were washed three times by magnetic separation, removal of supernatant and resuspension in 150 μl TNP-40 wash buffer. Subsequent to final wash, beads were re-suspended in 150 μl chilled LB and used to inoculate fresh E. *coli* cultures (400 μl at OD600 = 0.5) for *in vivo* amplification. Lysates were clarified by centrifugation at 3,000g (at 4 degrees) and 150 μl removed/stored for NGS library prep or additional rounds of IP.

### NGS Library Prep

Sequencing libraries were prepared from lysates with a single PCR reaction using multiplexing (MP) primers as follows:

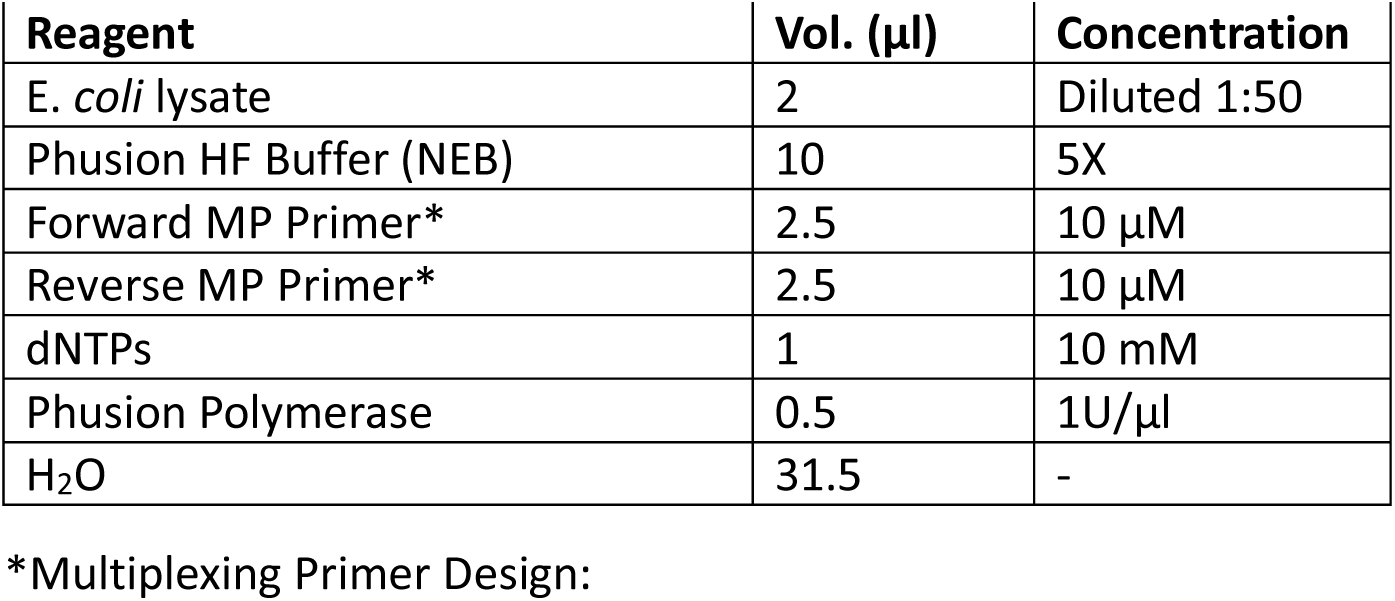

| Reagent | Vol. (µl) | Concentration |
| --- | --- | --- |
| *E. coli* lysate | 2 | Diluted 1:50 |
| Phusion HF Buffer (NEB) | 10 | 5X |
| Forward MP Primer* | 2.5 | 10 µM |
| Reverse MP Primer* | 2.5 | 10 µM |
| dNTPs | 1 | 10 mM |
| Phusion Polymerase | 0.5 | 1U/µl |
| H<sub>2</sub>O | 31.5 | - |

**\*Multiplexing Primer Design:**

Forward:

AATGATACGGCGACCACCGAGATCTACAC[NNNNNNN]GGAGCTGTCGTATTCCAGTCAGGTGTGATGCTC

NNNNNNN = i5 index

Reverse:

CAAGCAGAAGACGGCATACGAGAT[NNNNNNN]GGTTAACTAGTTACTCGAGTGCGGCCGCAAGCNNNNNNN = i7 index

Thermocycling conditions:

| Temperature (°C) | Time (mm:ss) | # cycles |
| --- | --- | --- |
| 95 | 02:00 | 1 |
| 95 | 00:30 | 5 |
| 63 | 00:30 |  |
| 72 | 01:00 |  |
| 95 | 00:30 | 5 |
| 60 | 00:30 |  |
| 72 | 01:00 |  |
| 95 | 00:30 | 10 |
| 58 | 00:30 |  |
| 72 | 01:00 |  |
| 72 | 02:00 | 1 |
| 4 | hold | - |

Libraries were cleaned and size selected by Ampure XP magnetic beads (0.8X volume, 40 μl) (Beckman Coulter, Pasadena, CA), eluted in 20 μl of nuclease-free water and quantified by Qubit (Thermo Fisher, Waltham MA) dsDNA high sensitivity fluorimeter. Sequencing was performed on either an Illumina HiSeq 2500 or 4000 using custom indexing and sequencing primers:

> Read 1 sequencing primer: TCCTGGAGCCATCCGCAGTTCGAGAAA
>
> Read 2 sequencing primer: AAGCTTCTTATCATCGTCGTCCTTGTAGTC
>
> i7 indexing primer: GGCCGCACTCGAGTAACTAGTTAACC
>
> i5 indexing primer: CACCTGACTGGAATACGACAGCTCC

### Bioinformatic Analysis

Reads were quality filtered, paired-end reconciled (PEAR v0.9.8),^50^ and aligned to a reference database of the full library (bowtie2 v2.3.1).^51^ Sam files were parsed using a suite of in-house analysis tools (Python/Pandas) and individual phage counts were normalized to reads per 100k (rpK) by dividing by the sum of counts and multiplying by 100,000. Peptide-level fold changes were calculated by dividing individual peptide rpK values with their expected abundance (mean rpK in mock IPs). For peptides never observed in any control IP, fold changes were calculated using the median rpK for all peptides derived from that gene in the mock IP.

We identified in our control IP’s a set of the 100 most abundant peptides that also have a standard deviation less than their mean. This “internal control” set represented the most abundant and consistent phage carried along specifically by the protein-AG beads or other reagents, in absence of any antibody. We calculate a sample-specific scaling factor, defined as the ratio of median abundances (rpK) of these 100 peptides in our controls to their abundance in the given sample. Fold change values are the multiplied by their sample-specific scaling factor. These top 100 peptide sequences are available at https://github.com/derisilab-ucsf/PhIP-PND-2018.

To account for the bias introduced in the fold-change calculation stemming from under-represented peptides in the mock IP having inflated fold-changes despite only modest representation in an experimental IP, we normalize, within each experimental batch (defined as all samples on a given 96-well plate - never less than 50 samples), fold-change values using a correction factor defined as the solution to a linear regression fit to the fold-change values as a function of their mean log2(rpK) in AG-only controls (Supplementary Figure 1). Importantly, fold-change values are only corrected when the correction factor is positive (expected fold-change given the abundance in mock IPs is positive). We fit a normal distribution to the scaled and normalized values and assign p-values by evaluating the survival function (1-CDF) at a given fold-change (Supplemental figure 1D). To account for multiple tests, we calculate an adjusted p-value (p_adj) using the Benjamini-Hotchberg procedure, enforcing a false discovery rate (FDR) of 0.05. Peptides were reported as significant as those with a positive fold-change with a p_adj <= 0.05 in both replicates.

### Design of Hu Motif Mutational Scanning Library

Degenerate 112 nt oligos were designed encoding 24 amino acids (PLHHQAQRFRLDNLLNMAYGVKRLMKL) spanning the 17-residue motif (QAQRFRLDNLLNMAYGVK) identified in the PhIP-Seq data. A unique oligo containing a degenerate NNN codon at each position was ordered/synthesized (IDT, San Diego, CA). Sequences were flanked with relevant restriction sites (EcoRI, HindIII) to enable in-frame cloning. Oligos were pooled in equimolar ratios and cloned following the manufacturer’s protocol in the T7 Select Cloning Manual. Phage libraries were subjected to 3 rounds of immunoprecipitation using patient antibodies (or AG beads alone) following the standard PhIP-Seq protocol described above. Final phage populations were sequenced with an average of ~2M 150 base paired-end reads on an Illumina MiSeq. Reads were quality filtered, reconciled/merged (PEAR),^50^ and translated to amino acid sequences. At each position, the antibody binding preference was defined as the log2(fold-change) of each amino acid substitution versus AG-only controls and relative to the endogenous/reference residue, quantified accordingly:

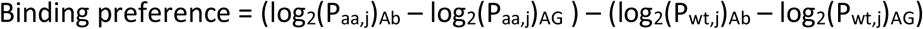

where:

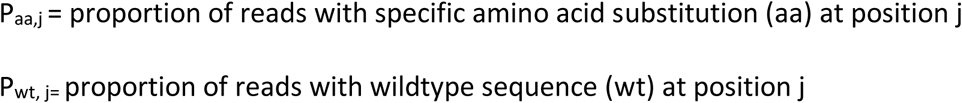

The mutational tolerance was then defined as the mean binding preference at each position.

### ELAVL4 amplicon sequencing from bulk thymus and sorted human TECs

Sorted TECs were isolated from human thymus obtained from 18- to 22-gestational-week specimens under the guidelines of the University of California San Francisco Committee on Human Research. Tissue was washed, cut into small pieces using scissors, and gently mashed using the back of a syringe to extract thymocytes. To isolate TECs, remaining tissue pieces were digested at 37 °C using medium containing 100 μg/ml DNase I (Roche, Belmont, CA) and 100 μg/ml Liberase™ (Sigma-Aldrich, St. Louis, MO) in RPMI. Fragments were triturated through a 5-ml pipette after 6 and 12 min to mechanically aid digestion. At 12 min, tubes were spun briefly to pellet undigested fragments and the supernatant was discarded. Fresh digestion medium was added to remaining fragments and the digestion was repeated using a glass Pasteur pipette for trituration. Supernatant from this second round of digestion was also discarded. A third round of enzymatic digestion was performed using digestion medium supplemented with trypsin–EDTA for a final concentration of 0.05%. Remaining thymic fragments were digested for another 30 min or until a single cell suspension was obtained. The cells were moved to cold MACS buffer (0.5% BSA, 2 mM EDTA in PBS) to stop the enzymatic digestion. Following digestion, TECs were enriched by density centrifugation over a three-layer Percoll gradient with specific gravities of 1.115, 1.065 and 1.0. Stromal cells isolated from the Percoll-light fraction (between the 1.065 and 1.0 layers) were washed in MACS buffer. For surface staining, cells were blocked using a human Fc Receptor Binding Inhibitor Antibody (eBioscience) and incubated on ice for 30 min with the following antibodies:

> PE anti-human Epcam (HEA-125) (Miltenyi 130-091-253) - dilution 1:50
>
> Alexa Fluor 488 anti-human CD45 (HI30) (BioLegend 304017) - dilution 1:100

Epcam+ CD45-DAPI-cells were sorted using a FACS AriaII (BD Biosciences, San Jose, CA) directly into TRIzol LS (ThermoFisher Scientific, South San Francisco, CA). The aqueous phase was extracted using the manufacturer’s instructions followed by RNA isolation using the RNeasy micro kit (QIAGEN, Redwood City, CA). RNA was reverse transcribed using iScript cDNA synthesis kit (Bio-Rad, Emeryville, CA).

RNA samples from whole fetal and adult human thymus were purchased from Agilent (Santa Clara, CA) and Clontech (Mountain View, CA). RNA was reverse transcribed using iScript cDNA synthesis kit (Bio-Rad, Emeryville, CA).

Amplicon libraries spanning the relevant exons were generated using the following primers:

> ELAVL4_fwd: GAACCGATTACTGTGAAGTTTGCCAAC
>
> ELAVL4_rev: GTAGACAAAGATGCACCACCCAGTTC

PCR products were purified using the QIAGEN Gel Extraction kit and library construction was conducted on the quantified DNA using the NEBNext^®^ Ultra™ DNA Library Prep Kit (New England Biolabs, Ipswich, MA). The final library was sequenced using a MiSeq with single-end, 300-base reads.

### Motif Discovery

The redundant and overlapping nature of our library design presents some potentially confounding issues with existing motif discovery software packages leveraging hidden Markov models (HMMs) to identify significantly represented subsequences and motifs in a collection of sequences or strings. Highly redundant and replicate sequences as an input dataset confound or overwhelm the motif finding algorithms, often resulting in the most significant motifs reported as those in the overlapping regions of adjacent peptides. To counter this, we manually curated patient results to eliminate sequences with >50% identity. We then iteratively searched each set of results for peptides with 10 or more identical sequential residues and chose as a representative sequence, that with the higher enrichment score (lowest adjusted p-value). These curated datasets were examined by the researchers before being analyzed for motifs by GLAM2, part of the meme-suite software package.^52^

### Clinical PND Antibody Confirmation

Confirmation of anti-Yo and anti-Hu activity in patient samples was performed in the Mayo Clinic Neuroimmunology Laboratory with CLIA and New York State approved methodology. Briefly, patient samples (serum/CSF) were tested on a mouse composite tissue slide with an indirect immunofluorescence assay for IgG binding corresponding to the anti-Hu or anti-Yo pattern, as previously described.^29^ The results were confirmed with western blot analysis on rat cerebellum preparations.

---

**SUPPLEMENTAL FIGURE 1:**
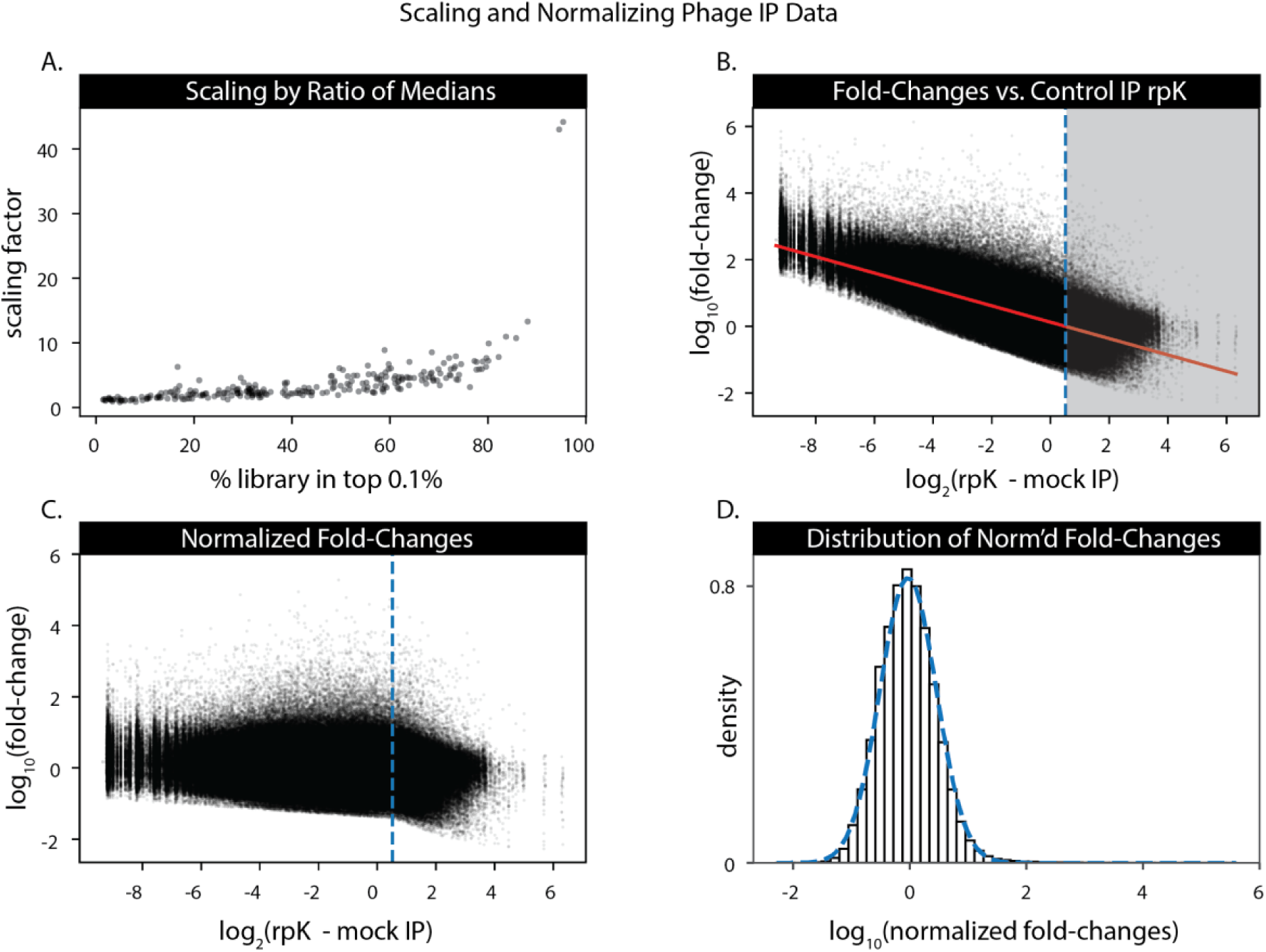
Development of a scaling and normalization schema for peptide fold-change values. A scaling factor, defined as the ratio of median rpK values of the top 100 mock IP AG bead-binding phage (see Methods), was multiplied by the normalized peptide count for a given experimental sample. A.) Plotting the scaling factor vs the proportion of an experimental sample’s final phage population present at or above the top 0.1% of rpK shows that samples dominated by a phage scale more dramatically. B.) Linear regression of scaled fold-change values to peptide representation in mock IPs. Intuitively, peptides with low abundance in the control IPs have higher calculated fold changes despite low rpK in the final population. Fold-change values were normalized by subtracting the solution to the fitted line when the expected fold-change was greater than 0. The resulting shifted/normalized fold changes (C.) are centered around 0 and normally distributed (D).

**SUPPLEMENTAL FIGURE 2:**
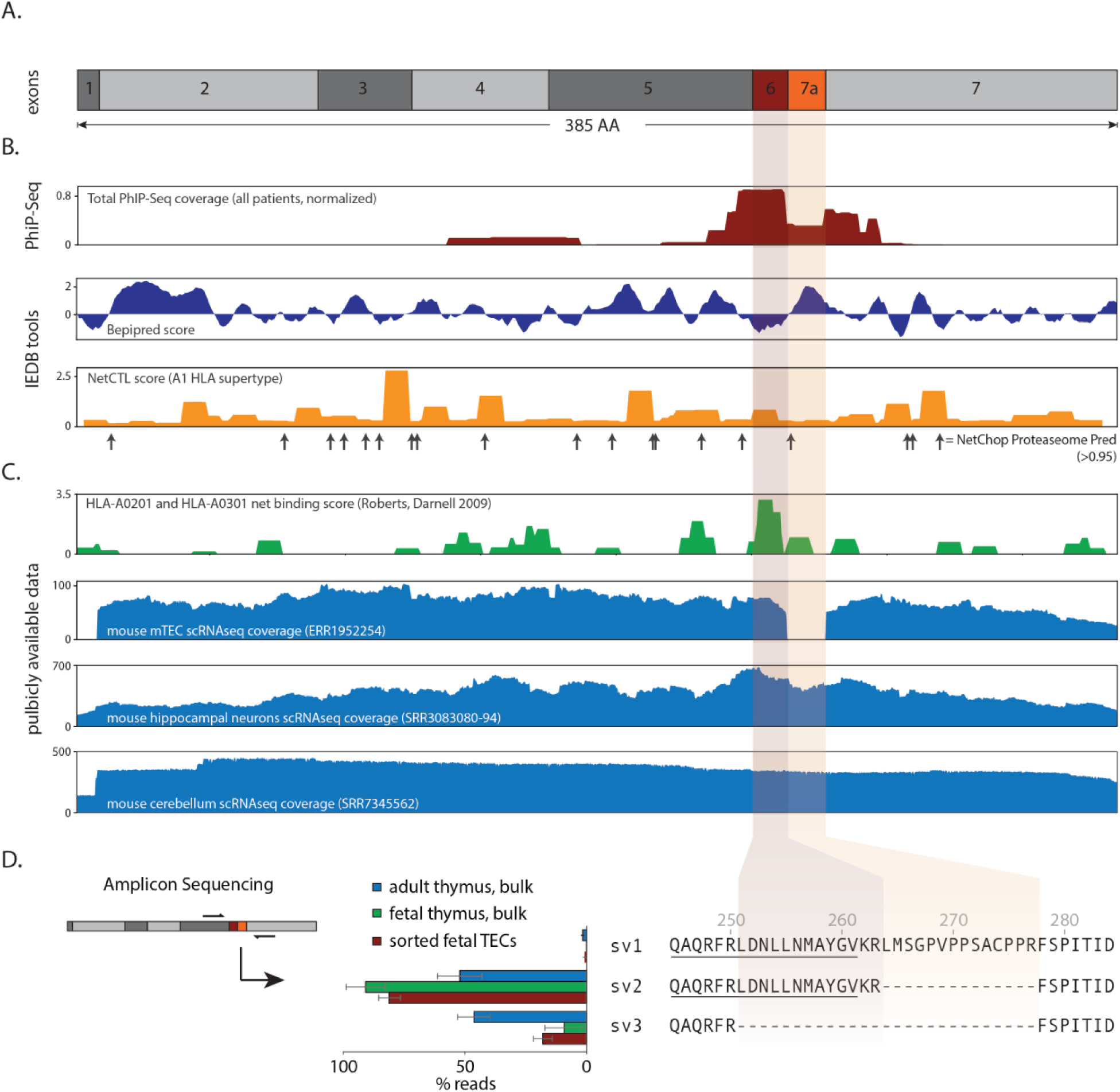
PhIP-Seq Identifies short exon in ELAVL4 excluded from expression in thymus. A.) Exon map of ELAVL4. B.) Normalized PhIP-Seq coverage from all positive patients shows preferential binding of antibodies to exon 6. Exon 6 has low predicted B cell antigenicity but high prediction of MHC processing and presentation by IEDB tools.^53^ C.) Exon 6 contains top scoring MHC-I binding prediction score defined by Roberts, et al.^33^ Analysis of publicly available datasets reveals exon 7a is excluded from transcripts in murine mTECs but included in mouse hippocampal and cerebellar neuron scRNAseq datasets. D.) Amplicon sequencing of human TEC cDNA corroborates exclusion of exon 7a in the thymus.

**SUPPLEMENTAL TABLE 1:** Preferred E. *coli* codons used in oligonucleotide library

| Amino Acid | Codon | Amino Acid | Codon |
| --- | --- | --- | --- |
| A | GCG | L | CTG |
| C | TGC | N | AAC |
| B | GAT | Q | CAG |
| E | GAA | P | CCG |
| D | GAT | S | AGC |
| G | GGC | R | CGT |
| F | TTT | U | TGC |
| I | ATT | T | ACC |
| H | CAT | W | TGG |
| K | AAA | V | GTG |
| J | CTG | Y | TAT |
| M | ATG | X | GCT |
|  |  | Z | GAA |

